# Defined extracellular ionic solutions to study and manipulate the cellular resting membrane potential

**DOI:** 10.1101/785444

**Authors:** Mattia Bonzanni, Samantha L. Payne, Myriam Adelfio, David L. Kaplan, Michael Levin, Madeleine J. Oudin

**Affiliations:** Department of Biomedical Engineering, Tufts University, Medford, Massachusetts; Allen Discovery Center, Tufts University, Medford, Massachusetts

**Keywords:** bioelectricity, ionic solutions, resting membrane potential, non-excitable cells, electrophysiology

## Abstract

All cells possess an electric potential across their plasma membranes. While familiar in the context of excitable cells such as neurons, healthy non-excitable cells are also able to generate and receive bioelectric signals. The cellular resting membrane potential (RMP) regulates many factors in cell homeostasis, such as cell proliferation, differentiation and apoptosis. It is therefore critical to develop simple strategies to measure, manipulate and characterize this feature. Current studies to evaluate RMP rely on the patch clamp approach, which is technically challenging, low-throughput and not widely available to the scientific community. Here, we present a relatively simple methodology to functionally study the role of RMP in non-excitable cells by modulating it pharmacologically, and using a voltage-sensitive dye to characterize the contribution of individual ions to the RMP. Specifically, we define protocols for using extracellular solutions in which permeable ions (Na^+^, Cl^−^ and K^+^) are substituted with non-permeable ions (N-Methyl-D-glucamine (NMDG), gluconate, choline, SO_4_^2−^) to study and manipulate RMP *in vitro*. The resulting RMP modifications were assessed with both patch clamp and a voltage sensitive dye. Using an epithelial and cancer cell line, we demonstrate that the proposed ionic solutions can determine the relative contribution of ionic species in setting the RMP and be used to actively and selectively modify the RMP. The proposed method is simple and reproducible and will make the study of bioelectricity more readily available to the cell biology community by enabling functional modulation of RMP in most cellular assays.

**Author Disclosure Statements:** No competing financial interests exist.

## Introduction

The uneven distribution of ions and their selective permeability across the plasma membrane leads to a difference in electric potential known as the membrane potential. Although commonly thought of in terms of electrically excitable cells such as neurons, every cell of the body generates a membrane potential. The resting membrane potential (RMP) represents the voltage at which the net ionic currents across the membrane are zero (Hille, 2001). When the membrane potential becomes more negative or positive than the RMP, the cell is said to be hyperpolarized or depolarized, respectively. The RMP is recognized as a key regulator of many cellular phenomena, such as proliferation, morphogenesis, migration, and differentiation, as well as morphogenesis in development and regeneration (Adams and Levin, 2013, Levin, 2014, McLaughlin and Levin, 2018, Mathews and Levin, 2018, Abdul Kadir et al., 2018). For example, it has been demonstrated that cells must undergo specific stages of depolarization and hyperpolarization to pass through the checkpoints of the cell cycle (Sachs et al., 1974). Membrane potential is also becoming increasingly linked to a wide variety of pathophysiological states, such as cancer (Yang and Brackenbury, 2013) and developmental disorders (Abdul Kadir et al., 2018), highlighting its role as a master controller of basic cell functions. Due to the biological importance of the RMP, it is necessary to define methods by which to experimentally study, manipulate and quantify it in cellular assays.

The RMP is determined by two main factors: 1) the membrane permeability and 2) concentration gradient of the most abundant permeable ions, Na^+^, K^+^, and Cl^−^ (Fig 1A). the voltage Goldman-Hodgkin Katz (GHK) equation is the simplest and most widely used method to calculate the RMP. The GHK equation describes the relationship between the RMP of a cell and variables such as temperature, ionic permeability and composition of the dominant monovalent ions of a cell: K^+^, Na^+^ and Cl^−^ (Fig 1B). Quantifying the contribution of each ion to the RMP can reveal insights into the cell biological mechanisms driving the RMP and the effects of changes in RMP within the cell (Fig 1C). The RMP is set and controlled by a repertoire of ion channels, pumps, transporters and junction proteins, whose activities are usually regulated by voltage, ligands, or both (Hille, 2001). The multifaceted time-, voltage-, spatial- and ligand-dependent activity and localization of these proteins allows the selective movement of ions across the membrane. As a consequence, the RMP itself varies in a cell specific, time- and context-dependent fashion, as previously summarized elsewhere (Levin et al., 2017), making it difficult to accurately capture using the GHK equation alone. Standardizing protocols to investigate the contribution of individual ions to the RMP and the effects of changes in RMP cell behavior are crucial to better understand the role of bioelectricity in cell biology. Due to the complex nature of the RMP, it is important to have a relatively easy, standardized and reproducible method to identify the major ionic players that set and control the RMP in different cell types, over time, or across different pathophysiological conditions.

**Figure 1:**
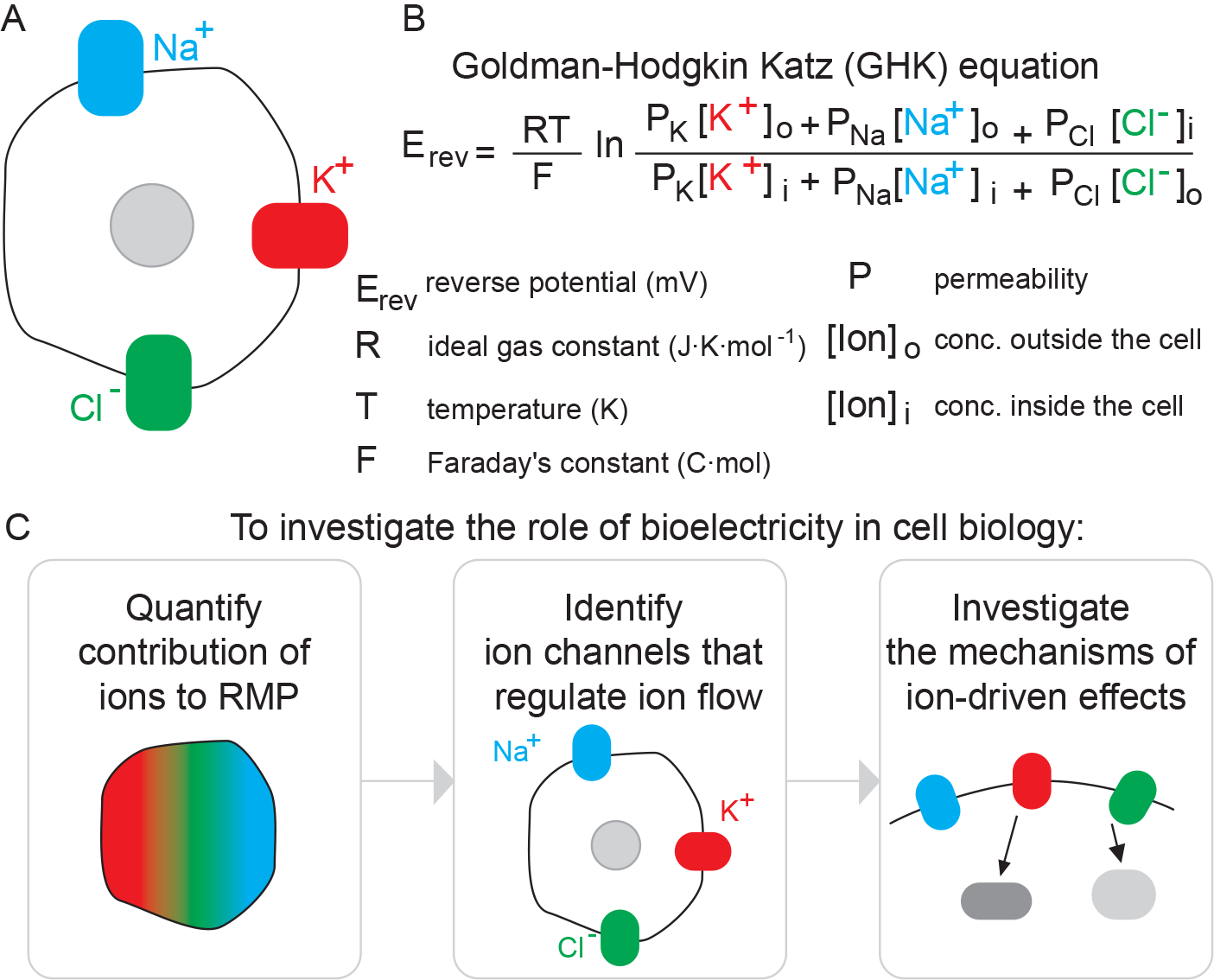
Investigating ion-driven effects on cellular membrane potential. **(A)** The RMP is mainly determined by the membrane permeability and concentration gradient of permeable ions Na^+^, K^+^, and Cl^−^. **(B)** The GHK equation describes the relationship between the RMP and other variables, including ion permeability. **(C)** Investigating the role of bioelectricity in a cellular process involves first identifying which ions are important in setting the RMP, next which channels regulate the flow of these ions, and finally probing the downstream signaling pathways of these ion-driven effects.

Current methods to evaluate RMP of a cell rely largely on patch clamp, a method which is technically challenging, low-throughput and not widely available to the scientific community. More recently, voltage-sensitive dyes have been developed which change their spectral properties in response to voltage changes. Although originally designed for measurements of the firing activity of neurons, slow membrane potential dyes such as Bis-(1,3-Dibutylbarbituric Acid)Trimethine Oxonol (DiBAC) are emerging as a method to provide optical quantification of the membrane potential in non-excitable cell types (Miller, 2016, Sundelacruz et al., 2008, Roger et al., 2007, Adams and Levin, 2012a, Oviedo et al., 2008). To understand the contribution of individual ions, most researchers rely on genetic or pharmacological tools to inhibit the activity or expression of individual ion channels. However, there are many different types of channels for a single ion and the function or activity of a single ion channel is not sufficient to capture the overall contribution of that ion to the RMP. Approaches such as RNAseq, immunohistochemistry and Western blot can potentially identify the membrane protein composition, but the inherently large number of ion-related proteins, and fluctuations in their expression and activity make it difficult to capture the dynamic nature of this phenomenon using these techniques alone. Moreover, it is important to emphasize that the RMP is not merely a function of the expression profile of ion channels, pumps, transporters and junction proteins or their individual activity. Indeed, the dynamic interaction between proteins (i.e. calcium channels and calcium-activated potassium channels) and with the cell microenvironment (i.e. actin, lipid rafts, caveolae, electric field) drastically influence the final RMP output.

To address the need for a simple and replicable technique for functional RMP analysis, we present a roadmap to study the role of RMP in a physiological process and to approximate the contribution of individual ions to the RMP. First, we describe a method to generate a calibration curve with voltage sensitive dyes to limit the need for electrophysiology. Second, we demonstrate the use of extracellular solutions to calculate the effect of changes in ion concentration on RMP. Finally, we define a series of extracellular solutions using non-permeable ions designed to isolate the effect of K^+^, Na^+^ and Cl^−^ individually on the RMP. The goal is to facilitate the study of bioelectricity in cell biology to a broader population of researchers.

## Material and Methods

### Cell culture

The MDA-MB-231 breast cancer cell line was obtained from ATCC and cultured in DMEM with 10% FBS (Hyclone). Cells were passaged regularly when they reached approximately 70% confluency and used between p5 and p15 for all experiments. For patch clamp, 1.5 × 10^5^ cells were seeded in an uncoated 2cm plastic dish 24 hours prior to the experiment. For voltage sensitive dye experiments, 1.0 × 10^4^ cells per well were seeded in an uncoated 96-well plastic plate 24 hours prior to the experiment. The renal proximal tube epithelial cell (RPTEC/TERT1) line was obtained from ATCC. RPTEC/TERT1 cells were cultured in DMEM/F-12 medium (Thermo Fisher), containing L-Glutamine and HEPES and supplemented with 25 ng/mL hydrocortisone (Sigma), 5 pM triiodo-L-thyronine (Sigma), 10 ng/mL recombinant human EGF (Thermo Fisher), 25 ng/mL prostaglandin E1 (Sigma), 3.5 μg/mL L-Ascorbic acid (Sigma), 1x ITS-G (Thermo Fisher) and 100 μg/mL geneticin (Thermo Fisher). For patch clamp and voltage sensitive dye measurements, RPTEC/TERT1 cells were seeded at 50% density into 35 mm dish and 12 well plate respectively and were grown for 3 days before starting the experiments. RPTEC/TERT1 cells up to p16 were used for all the experiments.

### Electrophysiology

Patch clamp experiments in the whole cell configuration were carried out at room temperature. Either MDA-MB-231 cells or RPTECs were superfused with different solutions as described in Table 2. Patch clamp pipettes had a resistance of 5-7 MΩ when filled with an intracellular-like solution containing (in mM): 130 K-Asp, 10 NaCl, 5 EGTA-KOH, 2 MgCl_2_, 2 CaCl_2_, 2 ATP (Na-salt), 5 creatine phosphate, 0.1 GTP, 10 Hepes-KOH; pH 7.2. Resting membrane potential (RMP) recordings were carried out in the I/0 configuration for at least 30 seconds or until steady state was reached. Neither series resistance compensation nor leak correction were applied: liquid junction potential correction was applied as previously reported(Neher, 1992).

### Voltage sensitive dye

Changes in RMP were measured using (Bis-(1,3-Dibutylbarbituric Acid)Trimethine Oxonol) (DiBAC; ThermoFischer). MDA-MB-231 cells, cultured in black clear-bottomed 96-well plates, were rinsed once with 200 μl of Fluorobrite medium (ThermoFischer) and then ionic solutions from Table 2 containing 1 μM DiBAC were added to cells and incubated in a cell incubator for 15 min to ensure dye distribution across the membrane. Assays were carried out at 37°C and 5% CO_2_ concentration in an incubation chamber fitted to a Keyence BZX700 fluorescence microscope. RPTEC/TERT1 cells, cultured in 12 well plates, were incubated with the solutions from Table 2 containing 50 nM of DiBAC for 30 minutes at 37°C and 5% CO_2_ concentration, before starting imaging (confocal microscope-Leica FLIM SP8). Changes in DiBAC fluorescence were measured at excitation and emission wavelengths of 488 and 520 nm, respectively and three regions of interest per well and three wells per solution for MDA-MB-231 cells and five regions of interest per well for RPTEC/TERT1 cells were imaged. DiBAC mean pixel intensity excluding background values was measured using ImageJ.

### Statistics

Patch clamp data were analyzed with Clampfit10 (Axon) and Origin Pro 9. Linear fitting was performed reporting the adjusted R^2^ value. Mean ± S.E.M. was reported from at least 2 different independent experiments for all the presented data. Either the number of cells or ROIs and the number of experiments were reported for each dataset.

## Results

### Generation of a standard calibration curve

The first step in the proposed method is the generation of a calibration curve to characterize the relationship between the optical reading from the voltage dye with the voltage reading by patch clamp within a cell line of interest.

The goal is to then use the change in fluorescence relative to the change in voltage obtained with the calibration curve measure the voltage difference between two conditions without the need for patch clamping for subsequent experiments. We used the DiBAC voltage-sensitive dye and quantified fluorescent signal for a field of view and patch clamp electrophysiology to measure the RMP of individual cells. To manipulate the RMP, we used a series of five electrophysiology solutions, which we describe here as calibration solutions (C1-C5), starting with ion concentrations approximating physiological conditions (C1) and increasing potassium and decreasing sodium concentrations (Fig 2). Potassium and sodium are two of the most abundant ions in the extracellular milieu and therefore were chosen to elicit a change in RMP. Altering the concentrations of both of these ions will impact the RMP in a dose-dependent manner to establish how changes in RMP correlate with changes in DiBAC fluorescence intensity.

**Figure 2:**
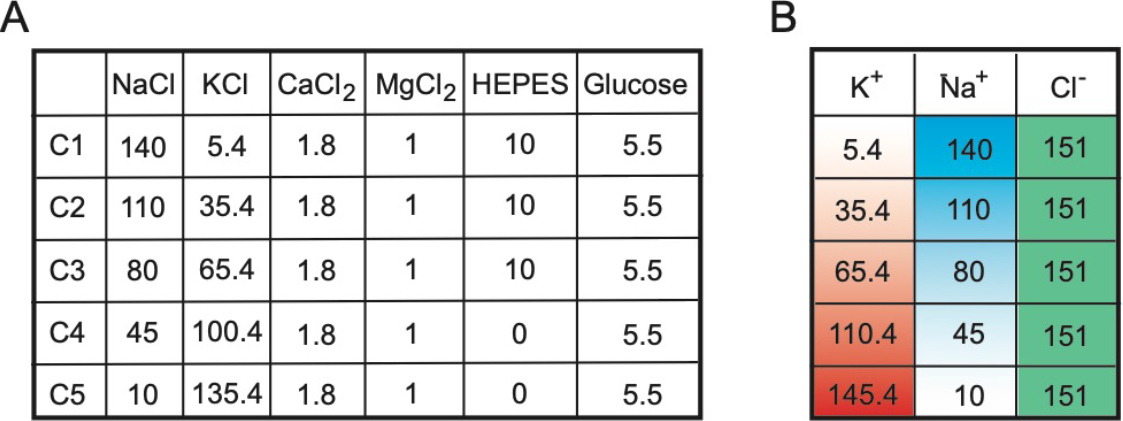
Ionic composition (in mM) of the extracellular solutions (pH=7.4) for generation of a fluorescence-voltage calibration curve. **(A)** Components of calibration solutions (in mM). **(B)** Change in K^+^. Na^+^ concentration with steady Cl^−^ concentration (in mM).

We first used renal proximal tubule epithelial cells (RPTECs) and exposed them to the five solutions. Increasing potassium and decreasing sodium concentration increased the DiBAC fluorescence signal (Fig 3A) as quantified in Fig 3B. The RMP values of individual cells were recorded with a patch clamp approach upon application of different solutions (Fig 3C). We combined voltage and fluorescence data, in which each point represents the voltage (x-axis) and optical (y-axis) readouts upon application of the same solution for a given cell line (Fig 3D) (Supplementary Data). For any given change in the fluorescence readout in two different solutions it is possible to calculate the relative voltage change:

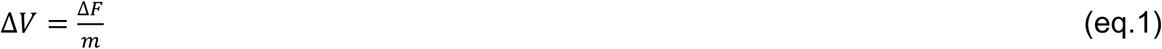

where Δ*V* is the voltage change, Δ*F* is the change in fluorescence, and *m* is the slope term from the linear fit. The slope of the resulting linear fit indicates the change in the fluorescence value per unit of voltage change (ΔF/ΔV), shown to be 1.76±0.19 F/mV for RPTECs.

**Figure 3:**
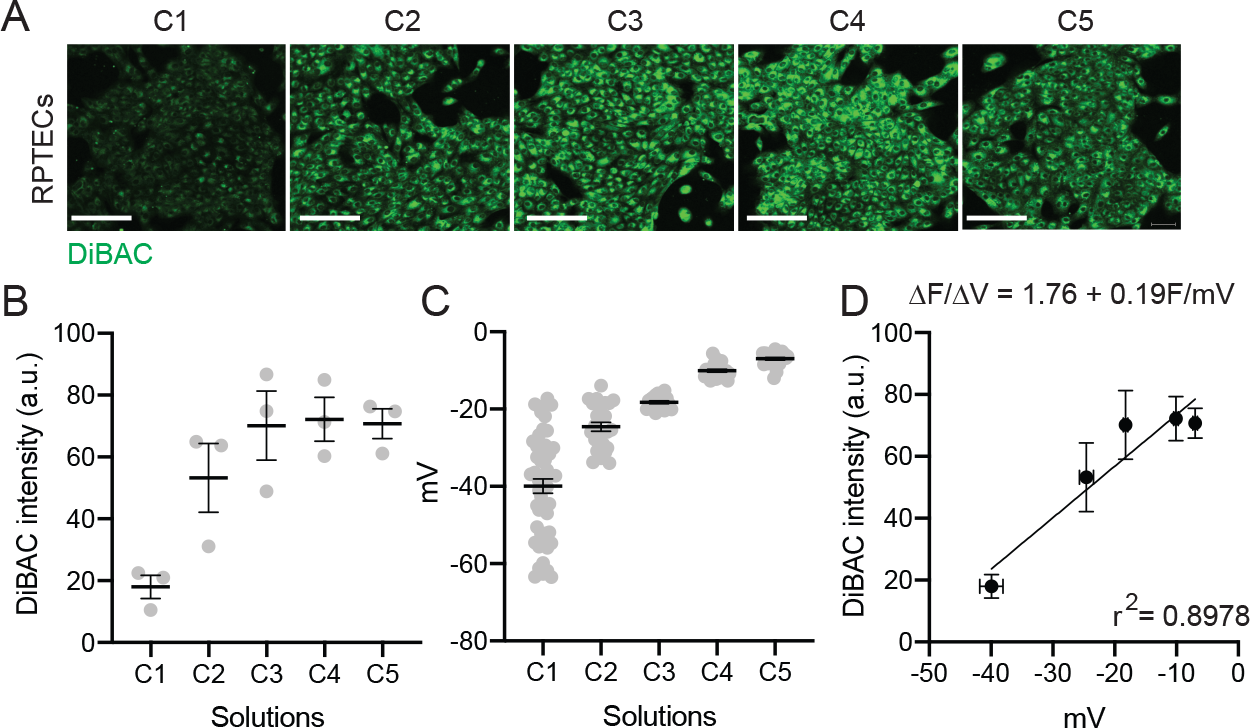
Generation of a fluorescence-voltage calibration curve in RPTECs using solutions C1-C5. **(A)** Representative images of DiBAC-stained cells in each calibration solution. **(B)** Mean DiBAC fluorescent intensity **(C)** Mean voltage RMP (mV) recorded by patch clamp **(D)** Fluorescence-voltage calibration curve with the change in fluorescence value per unit of voltage change expressed as the slope of the linear fit. Scale bar = 50um. Mean ± SEM.

### Calibration curves are cell-line specific

We next evaluated whether this method could be used to study bioelectric signaling in cancer cells, which are known to have a more depolarized RMP than epithelial cells(Williams et al., 2013). We used MDA-MB-231 triple-negative breast cancer cells and subjected them to solutions C1 to C5 outlined in Fig 2. Increasing potassium and decreasing sodium concentration increased DiBAC incorporation in the MDA-MB-231 cells (Fig 4A,B). Patch clamp electrophysiology of individual breast cancer cells reveals that the RMP increases in response to increasing K^+^ and decreasing Na^+^ in the calibration solutions (Fig 4C). We find a tight correlation between the RMP measured by patch clamp and DiBAC intensity in MDA-MB-231 cells, with a change in the fluorescence value per unit of voltage change (ΔF/ΔV) of 1.28±0.11 F/mV) (Fig 4D). These data demonstrate that calibration curves are cell-line specific and can be used to identify the differential effect of ion concentration on the RMP in healthy and diseased cells.

**Figure 4:**
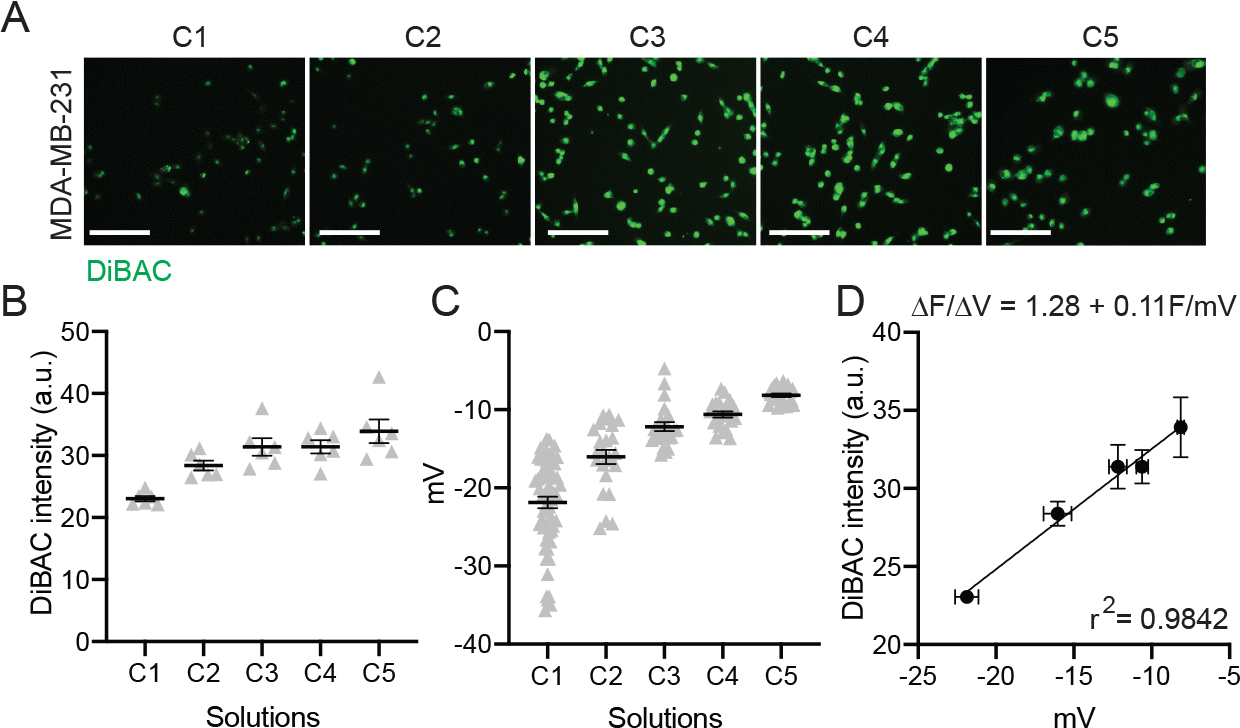
Generation of a fluorescence-voltage calibration curve in MDA-MB-231s using solutions C1-C5. **(A)** Representative images of DiBAC-stained cells in each calibration solution. **(B)** Mean DiBAC fluorescent intensity **(C)** Mean voltage RMP (mV) recorded by patch clamp **(D)** Fluorescence-voltage calibration curve with the change in fluorescence value per unit of voltage change expressed as the slope of the linear fit. Scale bar = 50um. Mean ± SEM. n=9 for C1 and n=3 independent experiments for C2-C5.

### Protocol for investigating the contribution of Na+, K+ and Cl− to membrane potential

We designed a simple methodology to identify the role of a given ion in setting the RMP by substituting each permeable ion with a cell membrane-impermeable ion and measuring the effect on the RMP. The Tyrode solution (Tyr) as the base starting point as it represents a widely used extracellular solution with a physiological distribution of ions. To determine the role of each ion, the Tyr solution is modified such that one permeable ion is replaced with an impermeable, while maintaining osmolarity and the electroneutrality in order to isolate the effect of a single ion species on establishing the RMP (Fig 5).

**Figure 5:**
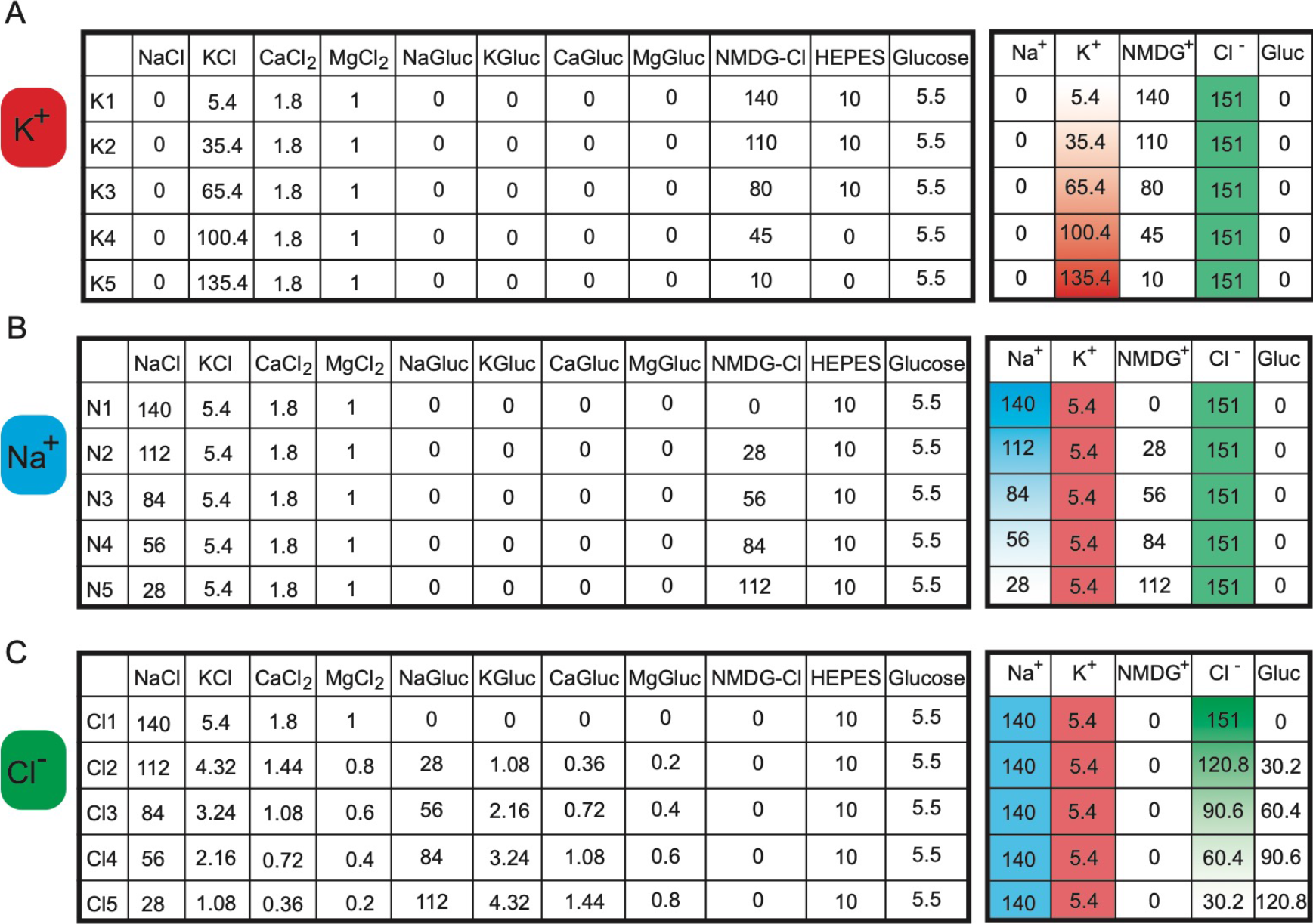
Ionic composition (in mM) of the extracellular solutions substituted with cell impermeable ions to isolate the contribution of each ion to the RMP. **(A)** K^+^ (K1-K5), **(B)** Na^+^ (N1-N5) and **(C)** Cl^−^ (Cl1-Cl5) solutions.

To study the role of K^+^, K^+^ can be replaced with N-Methyl-D-glucamine (NMDG) while removing Na^+^ (Fig 5A) To study the role of Na^+^, Na^+^ can be replaced with N-Methyl-D-glucamine while maintaining the K+ concentration at physiological level (Fig 5B). To study the role of Cl^−^, we can replace it with gluconate, while maintaining K^+^ and Na^+^ constant (Fig 5C). We recommend incremental ion reduction rather than the complete replacement of an ion to maintain cell viability over longer experimental time points and outline 5 solutions for each permeable ion to be used to identify their contribution to the RMP.

The effect of ion species on setting the RMP can be calculated by means of eq.1 comparing the RMP of cells in unmodified physiological conditions (Solution 1 from each table) with the RMP in modified solutions (Solutions 2-5) using the following formula:

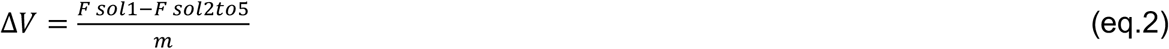

The resulting change in voltage induced by a specific solution is proportional to the importance of the reduced ion in setting the RMP.

## Discussion

The pathophysiological importance of membrane voltage is becoming increasingly recognized in many fields such as development (Bates, 2015, Levin and Martyniuk, 2018), regeneration (Zhao, 2009, McLaughlin and Levin, 2018), and cancer (Chernet and Levin, 2013, Onkal and Djamgoz, 2009) (Payne S.L., 2019). This increased recognition prompts the need for simple standardized methods to quantify and precisely manipulate the RMP and to identify the contribution of specific ions to the RMP. Indeed, interlaboratory variability due to the use of different experimental conditions (mainly different ionic solutions) accounts for up to 43% of the study-to-study variance in electrophysiological measurements in mammalian neurons (Tebaykin et al., 2018). Traditionally, measurement of membrane voltage has been confined to the study of fast fluctuations in excitable cells; however, the slow fluctuation of membrane voltage also play many crucial roles in non-excitable cells (Levin, 2014, Levin et al., 2017). The RMP, partially captured by the voltage Goldman-Hodgkin-Katz equation, is a function (among other parameters) of the uneven distribution and permeability of ions across the plasma membrane. To investigate the ionic permeability of a cell or the permeability of single channels to specific ions one approach is to manipulate the ionic composition of the extracellular solution (Hille, 2001, Chen et al., 2004, Knowles et al., 1983, Whittembury et al., 1961). However, to date, the lack of standardized extracellular and intracellular solutions makes the comparison of reported RMP values across studies more difficult. Moreover, the RMP is a non-thermodynamic equilibrium potential, which requires empirical evaluations rather than theoretical derivations (Hille, 2001). Here, we propose a roadmap to measure and manipulate the RMP which we anticipate can be used to study the role of RMP in cell biology. Specifically, we define a straightforward strategy (summarized in Fig 6) that involves the replacement of ions with non-permeable analogues to identify the contribution of these ions to the RMP.

**Figure 6:**
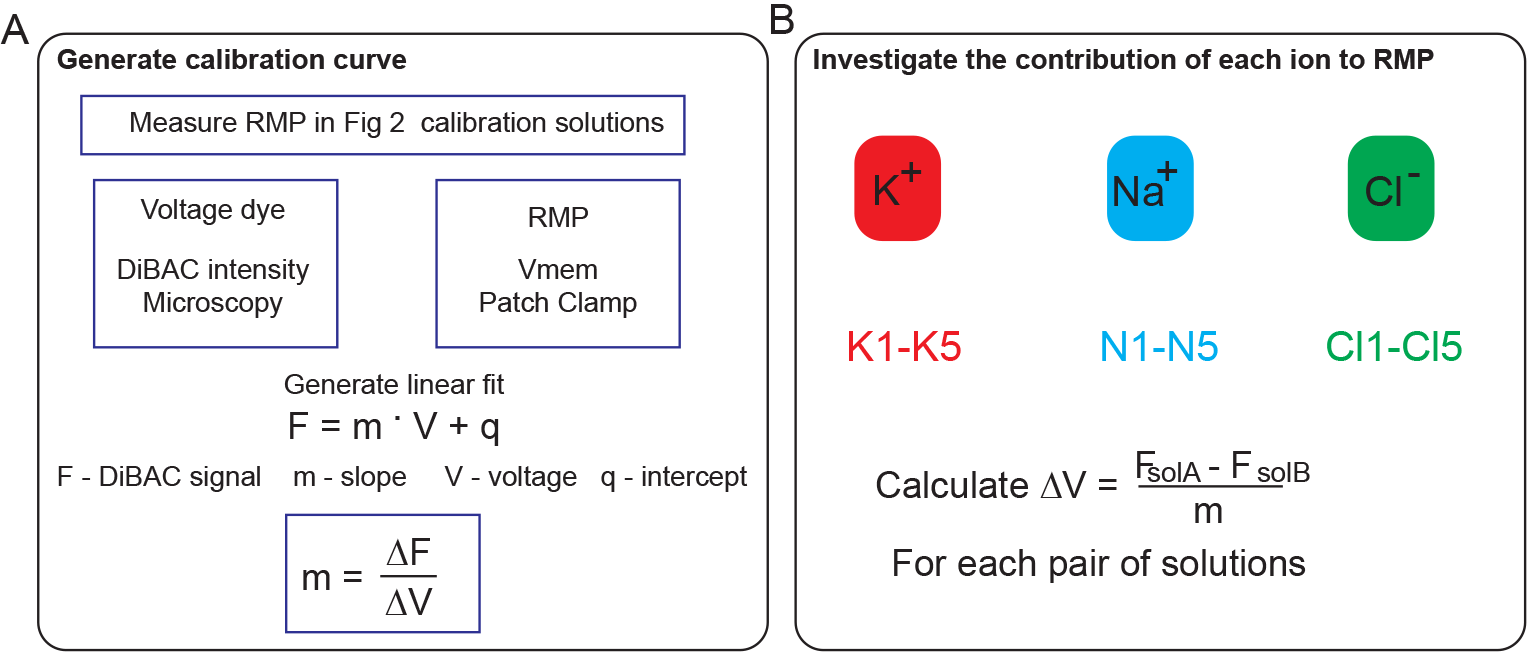
Proposed workflow for **(A)** the generation of a fluorescence-voltage calibration curve and **(B)** the determination of the contribution of cell-membrane permeable cations and anions to the RMP.

The first step in our proposed method is to generate a fluorescence-voltage calibration curve to obtain the variation of fluorescence for unitary variation of voltage, by combining patch clamp and voltage-sensitive dye optical approaches (Fig 6A). While patch clamp remains the gold standard for its precise and control, it requires specific equipment and expertise and involves membrane rupture. The use of voltage dyes increases the throughput of voltage studies and preserves the intracellular environment (Gerencser et al., 2012); it is also suitable for use in moving cells or whole tissues.

We recommend 5 solutions for the calibration, which provide a range in both K+ and Na+ concentration to cause a sufficiently wide change in RMP as a proof of concept. After the construction of the calibration curve for a given cell type, the optical readout will preserve the intracellular environment and can be used as a substitute for patch clamp analysis to measure a change in membrane potential, improving the throughput when testing different solutions and /or ion channel blockers. Using this method, we demonstrated that calibration curves are cell line-specific and can be obtained for both healthy and disease-state cell lines. The DiBAC dyes are well established for use in cells (Adams and Levin, 2012b, Yamada et al., 2001, Klapperstuck et al., 2009) and in vivo {Durant, 2019, 30799071;Adams, 2016, 26864374;Vandenberg, 2011, 21761475;Beane, 2011, 21276941;Pai, 2012, 22159581;Pai, 2018, WOS:000427007700007}, but theoretically any voltage-sensitive dye can be used to generate a fluorescence-voltage calibration curve. However, it is important to note that we cannot assign absolute voltages for any corresponding fluorescent value from the fluorescence-voltage-calibration curve since whole cell configuration of patch clamp accesses the cytosol, whereas DiBAC associates with the intact cell membrane. This difference prevents any direct relationship between single voltage values measured with the two different techniques, allowing only for a comparison between variations of fluorescence values per unit of voltage. Therefore, this technique is best suited for comparing biological conditions and understanding how different experimental conditions may impact RMP.

Second, we define a series of extracellular solutions that can be used to identify the contribution of individual ions to the RMP (Fig 6B). This technique works by changing the electromotive force of a specific ion, which will impact the RMP if this ion is involved in its establishment. This workflow provides a standardized method to easily screen for the most relevant ions in setting the RMP of a cell line which can then be complemented by approaches to target channel subtypes, such as pharmacological inhibition of channel activity. Unlike other methods which target a single channel subtype, by limiting the availability of a cell-permeable ion in the extracellular solution, we can assess the global contribution of that ion to the RMP in terms of all channels, pumps, and transporters. We demonstrate that this method works for both healthy and disease-state cell lines, and could be used to test ion permeability across time and different experimental conditions. Furthermore, by standardizing both ionic solutions and a workflow, results obtained by this method in different research environments can be directly compared.

There are a number of important considerations when using this method. The main caveat is the inability to control the intracellular ionic composition, limiting the parameters of the GHK equation that we can actively manipulate to the [Ion]_out_. Moreover, it is important to consider the nonlinear dynamic nature of the GHK equation; for example, the voltage dependence of the probability terms in the equation. This implies that the final RMP value is not just a function of the substituted ion(s), but also of the voltage-dependent change of every permeability variable. Furthermore, since it is possible that the cell is permeable to either NMDG (Wang et al., 2009, Harkat et al., 2017) or gluconate (Chen et al., 2004), the use of more than one osmolarity non-permeable cationic (choline (Knowles et al., 1983)) or anionic ion (i.e. SO_4_^2−^) is strongly recommended to confirm findings. Lastly, here we focused on K^+^, Na^+^, Cl^−^ but it is worth noting that other ions could, directly or indirectly, contribute to the membrane potential, such as Ca^2+^, Mg^2+^, HCO_3_^−^, and HPO_4_^−^. However, the same experimental design we have presented here can be used to study the contribution of these ions. As this method of RMP manipulation depends on changing the ion concentration and ions themselves are involved in other signaling pathways independent of membrane potential, it is crucial to test if the effect that we observe is ion- rather than purely voltage-dependent (Adams and Levin, 2013). The easy way is to empirically reach the same absolute value of RMP by manipulating different ions and determining if the recorded output is ion-invariant. In other words, if it is possible to reach the same RMP (either depolarizing or hyperpolarizing the initial membrane voltage) by manipulating different sets of ions without changing the cell output, this suggests that the cell output itself is ion-invariant and membrane voltage-dependent.

The use of these solutions can be complemented by other approaches. Combining this technique with targeted ion channel drugs (Kubo et al., 2005, Goldstein et al., 2005, Gutman et al., 2005, Wei et al., 2005, Clapham and Garbers, 2005, Clapham et al., 2005, Hofmann et al., 2005, Catterall et al., 2005b, Catterall et al., 2005a) could rule out the relative contribution of specific ion channels after narrowing the most relevant ion species and/or based on previous knowledge of mRNA/protein levels (the % of effect could be plotted against the range of concentrations of the tested drug). Moreover, we can take advantage of commonly used ionophores (i.e. valinomycin) to change the membrane potential of a cell; namely, we can increase the permeability of the cell to a single ion and observe the effects in our solution series. The RMP will thus coincide with the Nernst’s potential for that ion. In addition, we can also isolate and measure the pH-dependence of the RMP by preparing the Tyr solution with different pH values.

In the present work, we propose defined ionic solutions not only to study the role of ions in establishing the RMP, but most importantly to manipulate it. This method will be instrumental in revealing the role of ions and/or voltage in physiological (e.g. cellular migration, patterning, proliferation) or pathological processes (e.g. cancer growth and invasion, induction of fibrosis), across different cell types. As previously highlighted (Tebaykin et al., 2018), to reach a consensus of the composition of defined extracellular solution will also be instrumental to build new reliable, reproducible and solid RMP-related results to compare, across different cell types, conditions, and studies, the role of ions and membrane voltage.

